# Genome-wide analysis reveals genes mediating resistance to paraquat neurodegeneration in *Drosophila*

**DOI:** 10.1101/2025.04.02.646829

**Authors:** Stefanny Villalobos-Cantor, Alicia Arreola-Bustos, Ian Martin

**Affiliations:** Jungers Center for Neurosciences, Department of Neurology, Oregon Health and Science University, Portland, Oregon, USA; Department of Chemical Physiology and Biochemistry, Oregon Health and Science University, Portland, Oregon, USA

**Keywords:** Parkinson’s disease, Drosophila, paraquat, neurodegeneration, dopamine

## Abstract

Parkinson’s disease (PD) is thought to develop through a complex interplay of genetic and environmental factors. Epidemiological studies have linked exposure to certain pesticides such as paraquat with elevated PD risk, although how a person’s genetic makeup influences disease risk upon exposure remains unknown. Here, we used a genome-wide approach to uncover genes that play a role in resistance to paraquat-induced dopaminergic neurodegeneration in *Drosophila*. We developed a paraquat exposure model displaying delayed-onset dopaminergic (DA) neurodegeneration to recapitulate this aspect of human disease. We reveal that genetic background is a strong determinant of paraquat-induced DA neurodegeneration susceptibility across a series of nearly 200 fly strains called the *Drosophila* genetic reference panel (DGRP). Through unbiased genome-wide analysis and follow-up validation, we identify two candidate paraquat resistance genes, *luna* and *CG32264*. In gene-level studies, decreased expression of *luna* or *CG32264* is associated with paraquat-induced DA neuron loss while overexpression of either gene prevents neurodegeneration *in vivo*. The mammalian ortholog of *CG32264* is *Phactr2*, which has previously been linked to human idiopathic PD risk in several populations. Hence, our results reveal genes regulating paraquat-induced DA neuron loss that intersect with human PD risk variants, supporting the potential relevance of our findings to PD and underscoring a role for gene-environment interactions in pesticide-related DA neurodegeneration.

**ARTICLE SUMMARY:** Paraquat is a widely used herbicide linked to increased PD risk and to dopaminergic neurodegeneration in animal studies. Gene-environment interactions likely influence whether an individual exposed to paraquat eventually manifests PD and presents a major opportunity to yield insight into PD genetics. We developed a paraquat-induced neurodegeneration model in *Drosophila*, applied this model to nearly 200 fly strains belonging to the *Drosophila* Genetic Reference Panel and used a genome-wide association approach to identify candidate modifier genes of paraquat-induced dopamine neuron loss which we subsequently validated through RNAi and overexpression functional testing. Through this approach, we reveal two novel paraquat resistance genes, *luna* and *CG32264*. Strikingly, the putative mammalian ortholog of *CG32264* (*Phactr2*) was previously linked to human PD, supporting the potential relevance of our findings to human disease.

## INTRODUCTION

Genetic, environmental and aging factors converge to determine a person’s lifetime risk of developing Parkinson’s disease (PD), yet for most PD cases disease etiology remains unknown (1). Epidemiological studies have linked exposure to numerous types of environmental toxins with elevated PD risk, although pesticides have received the most attention due to concerns over the potential widespread impact of dietary, occupational and residential exposures to human health (1). Several pesticides including rotenone, paraquat, organochlorines and dithiocarbamates have been implicated in PD (2–4). A meta-study of 46 individual pesticide studies reported a summary PD risk ratio of 1.6 (95% confidence interval 1.4-1.9) for ever (versus never) pesticide exposure (5). A corollary of this risk metric is that most individuals exposed to pesticides do not develop PD, at least within the timeframe of these studies. While risk is likely dependent on the extent of pesticide exposure and on interaction with other environmental factors (6), it has been postulated that individual genetic architecture also influences the likelihood of manifesting disease via gene-environment interactions (7). A clear role for genetics in PD pathogenesis has emerged over the last few decades from the identification of pathogenic mutations within several genes that can cause familial PD (8). More recently, genome-wide association studies (GWAS) have revealed numerous loci harboring genetic variants associated with idiopathic PD risk (9), demonstrating that background genetic variation also plays a role in non-familial disease. Genes emerging from these GWAS have increased our understanding of genetic risk factors in idiopathic PD populations (9, 10) and genome-wide association for PD linked to pesticide exposure could in theory uncover important molecular mediators of dopaminergic neurodegeneration that pave the way for novel therapeutic strategies. Yet, such human studies are hindered by the challenge of obtaining a sufficiently large number of cases with accurate exposure assessment needed for the statistical power to detect genetic associations and additionally, the cost associated with genotyping those cases (4, 11).

Controlled animal models play a vital role in determining whether exposure to a given pesticide can cause PD-related neurodegeneration and can further be leveraged to gain mechanistic insight into disease etiology (6). Genetically tractable model organisms such as *Drosophila* can facilitate unbiased efforts to identify genes associated with environmental drivers of PD-related neurodegeneration such as pesticides (12). Animal studies can be conducted in tightly controlled environmental conditions, where pesticide concentration and exposure duration can be carefully regulated. In addition to the extensive conservation of gene and nervous system function shared between flies and vertebrates, the *Drosophila* brain contains dopamine neurons that are important in the control of movement (13). The vast array of sophisticated genetic tools available for *Drosophila* enables a rapid path for identifying disease-related genes that can be assessed for relevance in mammalian models (14). One such tool, the *Drosophila* Genetic Reference Panel (DGRP), is a set of ∼200 fly strains that have been inbred to homozygosity and fully sequenced, allowing the identification of genes associated with complex traits (15). The DGRP have previously been utilized to study neurodegenerative phenotypes associated with Aβ_42_, tau and mutant LRRK2 G2019S toxicity (16, 17). Here, we used the DGRP to uncover genes associated with paraquat-induced dopaminergic neurodegeneration. Paraquat, one of the most widely used herbicides in the world, crosses the blood-brain barrier and promotes DA neuron death in several animal models including *Drosophila*. Paraquat generates reactive oxygen species (ROS) via redox cycling and most animal studies to date have focused on the contribution of ROS and mitochondrial dysfunction as potential mechanistic drivers of neurodegeneration. Yet, the full scope of paraquat neurotoxicity and key mechanisms driving neurodegeneration remain unclear (18). Here, we reasoned that an unbiased identification of genes associated with paraquat-mediated DA neuron death may yield new insights into pathways mediating neurodegeneration. Our genome-wide association approach uncovered a number of candidate genes mediating paraquat resistance.

Through rigorous validation of candidate genes by RNAi and overexpression analyses, we report that *CG32264* and *luna* are associated with paraquat neurodegeneration. *luna* has not previously been linked to PD-related neurodegeneration, but *CG32264* is the *Drosophila* ortholog of *Phactr2*, a gene which has been linked to human idiopathic PD susceptibility. Collectively, our results reveal candidate modifier genes for paraquat-induced DA neuron loss that intersect with human idiopathic PD genes, supporting the contribution of gene-environment interactions to pesticide-mediated DA neurodegeneration.

## RESULTS

### Paraquat-induced DA neuron loss is divergent across genetic backgrounds

Epidemiological evidence indicates that PD associated with chronic paraquat exposure typically manifests years to decades after exposure (4). Towards modelling slow-onset DA neurodegeneration and avoiding potential mortality confounds associated with acute exposure to high neurotoxin doses, we sought a paraquat concentration that, when added to our standard fly food, would cause delayed-onset DA neuron loss and not cause major lethality. Exposure to food supplemented with 10 mM paraquat results in substantial DA neuron death after just 7 days of exposure in a prototypic DGRP strain, RAL-911 (Fig. 1A). Conversely, flies from this strain exhibit no immediate DA neuron loss upon exposure to 1 mM paraquat food for 7 days whereas they manifest significant DA neuron loss after being housed on standard fly food without paraquat for an additional 14 days immediately after paraquat exposure (Fig. 1B-C). This is consistent with a slow onset of neuronal death at the lower paraquat concentration. Flies exposed to 1 mM paraquat retain 80% survival at the end of the 21-day testing period suggesting that survival is not substantially compromised. We assessed food consumption rates to determine whether supplementation with paraquat at this concentration affects feeding behavior, treating dietary restriction as a potential confounder. Food intake, measured by ingestion of erioglaucine dye-containing food by *w^1118^* flies as described previously (19), is not affected by the presence of paraquat at this concentration in fly food (Fig. 1E), consistent with no effect on feeding behavior.

**Figure 1.**
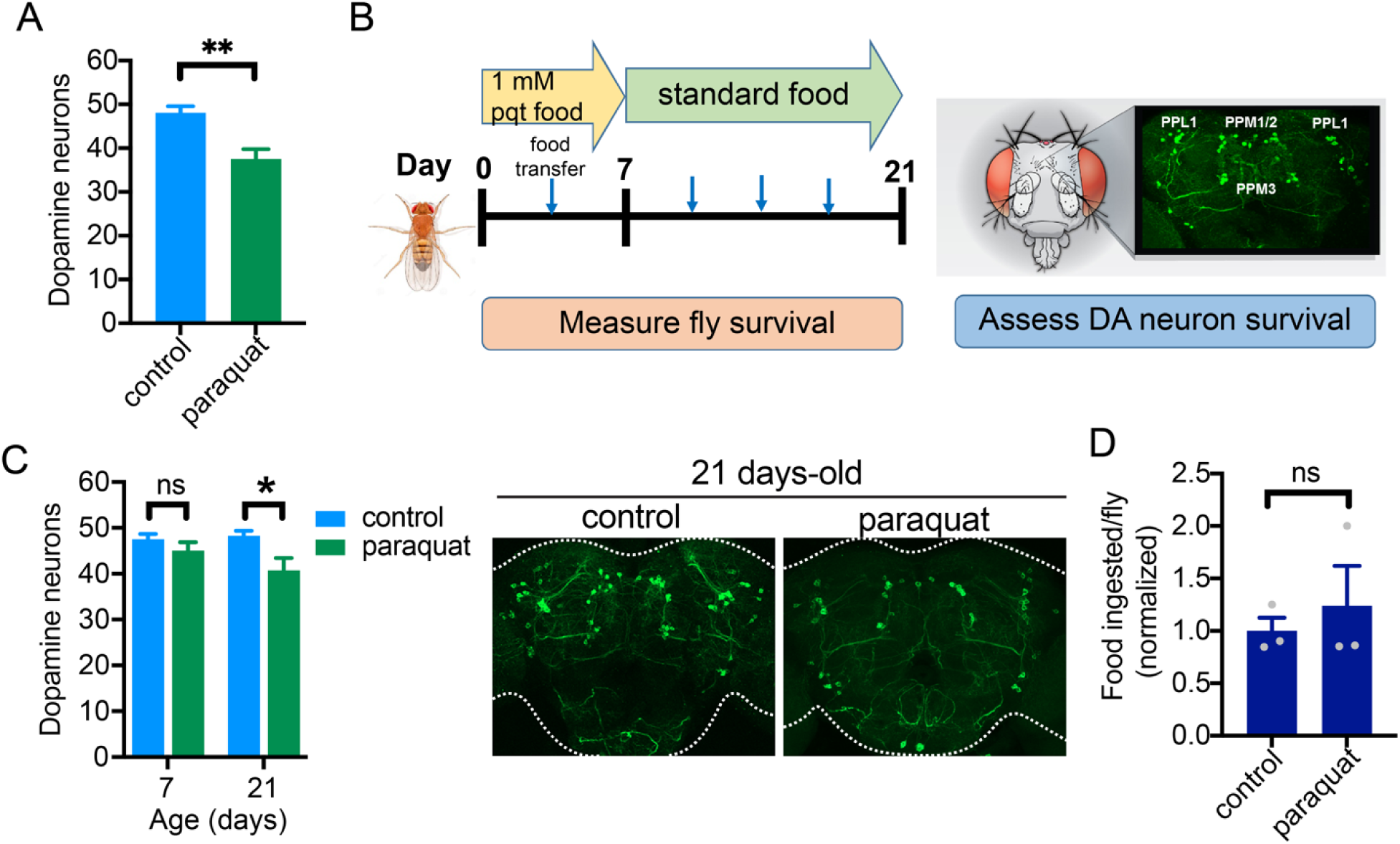
Slow-onset DA neurodegeneration in flies consuming paraquat. (A) housing flies on 10 mM paraquat-supplemented food for 7 days results in significant loss of DA neurons (Student’s t-test, *P* <0.01, *n* =12-15 brains). (B) scheme illustrating paraquat exposure and testing timeline (C) DA neuron viability in RAL-911 is not affected by paraquat exposure at day 7, but is at day 21 (individual 2-way ANOVAs). (D) Feeding behavior in *w^1118^*flies is not affected by paraquat food supplementation (Student’s t-test, n.s).

We applied this paraquat exposure model to 173 DGRP strains and identified substantial variation in DA neuron viability following paraquat exposure across strains (ANOVA, *P* = 6.09 x 10^-128^) (Fig. 2A and Table S1). Using individual brain DA neuron counts, we estimate a broad sense heritability for DA neuron survival on paraquat food of *H^2^* = 83.4% ±1.8%. Some strains exhibit near-maximal DA neuron numbers following paraquat exposure consistent with paraquat resistance, whereas others display fewer neurons, consistent with paraquat sensitivity (Fig. 2B). We considered the possibility that divergence in DA neuron numbers between strains could occur independently of paraquat, i.e., be developmental in origin or, since we assessed 3-week-old flies, that DA neuron integrity could be affected by aging. To probe these possibilities, we compared our measures of DA neuron viability with a prior study on the DGRP in which PPL1 DA neurons for most of the same strains were counted in 21-day-old flies not exposed to paraquat (Fig. 2C). We observe a weak Pearson correlation between our measures of PPL1 DA neuron viability following paraquat exposure and PPL1 DA counts at the same age derived from this prior study (20) (*r* =0.21, *P* <0.01, *n* =173), suggesting that neither strain-specific development nor aging are major sources of DA neuron phenotypic variation among the DGRP strains in our study. Since environmental differences between labs could affect the validity of this comparison, we aged a subset of 20 DGRP strains for 21d and assessed DA neuron viability to compare phenotypes across lines housed in the same laboratory either exposed to paraquat or unexposed.

**Figure 2.**
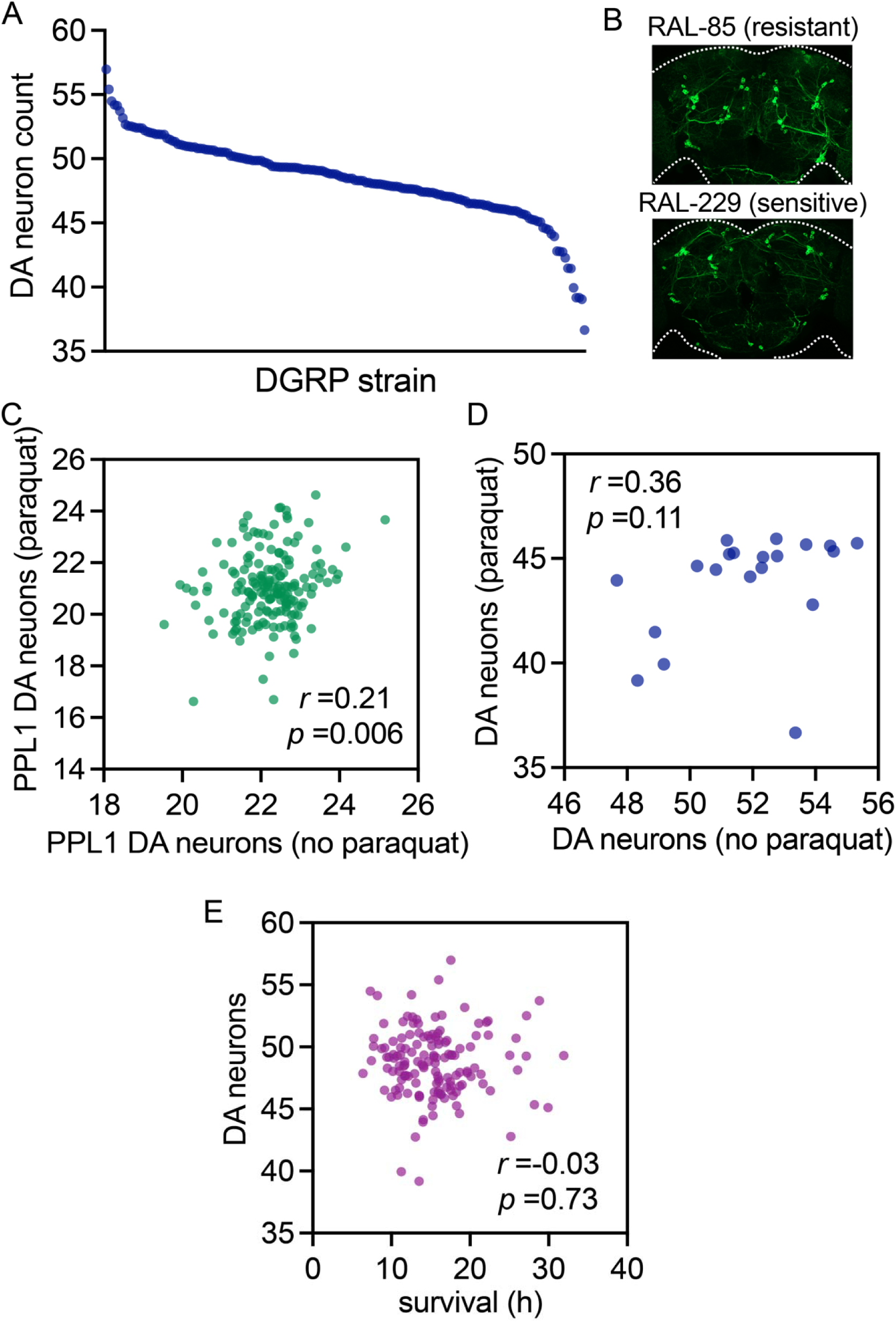
DA neuron viability following paraquat exposure diverges with natural genetic variation. (A) DA neuron viability in 173 DGRP backgrounds following 7d of paraquat exposure and 14d additional aging. DGRP background has a strong effect on DA neuron counts (*P* = 6.09 x 10^-128^). (B) examples of a paraquat-resistant (RAL-85) and a paraquat-sensitive (RAL-229) DGRP strain (C) weak correlation of PPL1 DA neuron viability between paraquat exposed DGRP strains (this study) and those aged without paraquat exposure for 21d (adapted from Davis *et al*. (Ref 20)). (D) weak correlation of DA neuron viability between paraquat exposed DGRP strains and those aged without paraquat exposure for 21d (E) no correlation of DA neuron viability following 1 mM paraquat exposure with survival of DGRP strains on 20 mM paraquat (adapted from Weber *et al*. (Ref (19)). Pearson *r* values indicated.

We chose 20 DGRP strains with paraquat-related DA neuron viability in the bottom 20^th^ percentile of all strains, as these strains exhibit the most dynamic range of DA neuron numbers (Fig. 2A). We find no significant correlation of DA neuron numbers between paraquat exposed and unexposed flies for these 20 strains (*r* =0.36, *P* =0.11), again supporting paraquat exposure as a major driver of DA neuron phenotypes across strains in our study. Additionally, all 20 DGRP lines tested possess more DA neurons in the absence of paraquat exposure (mean =7.6 ± 0.4 neurons) consistent with paraquat causing a loss of DA neurons and arguing against a developmental origin for reduced DA neuron numbers in these strains. Mean (± standard error) survival of all 173 lines housed on paraquat food is 97% ± 0.38% at the end of the 21-day testing period (Table S1), thus flies examined for DA neuron viability at the end of the 21d period are considered representative of their DGRP strain populations. Another prior study (21) examined DGRP strain survival on a relatively high dose of paraquat (20 mM) which induces rapid mortality with a mean survival time <35h for all strains tested. On comparing our paraquat DA neuron measures with strain-level paraquat survival we observe no correlation (*r* =-0.02, *P* =0.76, *n* =141), effectively uncoupling rapid-onset mortality of flies exposed to a relatively high dose paraquat from the DA neuron degeneration phenotype assessed in our study (Fig. 2D).

### Genome-wide association reveals candidate paraquat resistance genes

DA neuron counts for 173 DGRP strains were tested for association with genome-wide single nucleotide polymorphisms (SNPs) totaling ∼1.9 million. Candidate SNPs are filtered for MAF>0.05 and <30% missing genotypes. This approach is not sufficiently powered to identify SNPs with genome-wide significance that survive multiple testing correction but can effectively yield candidate gene associations suitable for further rigorous validation (17, 22). We identified 229 SNPs in 133 candidate genes associated with paraquat-mediated DA neuron death at a nominal P-value threshold of *P* <10^-5^ (Table S2) and 30 SNPs in 22 candidate genes at a more stringent threshold of *P* <10^-6^ (Table 1). These 30 SNPs are distributed throughout the fly genome (Fig. 3A) and pairwise linkage disequilibrium (LD) analysis indicates that they associate with paraquat-mediated DA neuron death as 24 independent loci (*r^2^* <0.5) consistent with a low impact of LD on the GWAS results (Fig. 3B). Functional annotations for candidate genes nominated by these SNPs encompass a broad range of biological processes including regulation of synaptic plasticity (*kek6*), retrograde vesicle-mediated transport (*zetaCOP*), DNA transcription, replication and demethylation (*luna*, *TH1*, *mei-41* and *tet*), translational regulation (*msi*) and actin cytoskeleton organization (*sif* and *CG32264*). Most candidate genes have putative human orthologs (Table 1).

**Figure 3.**
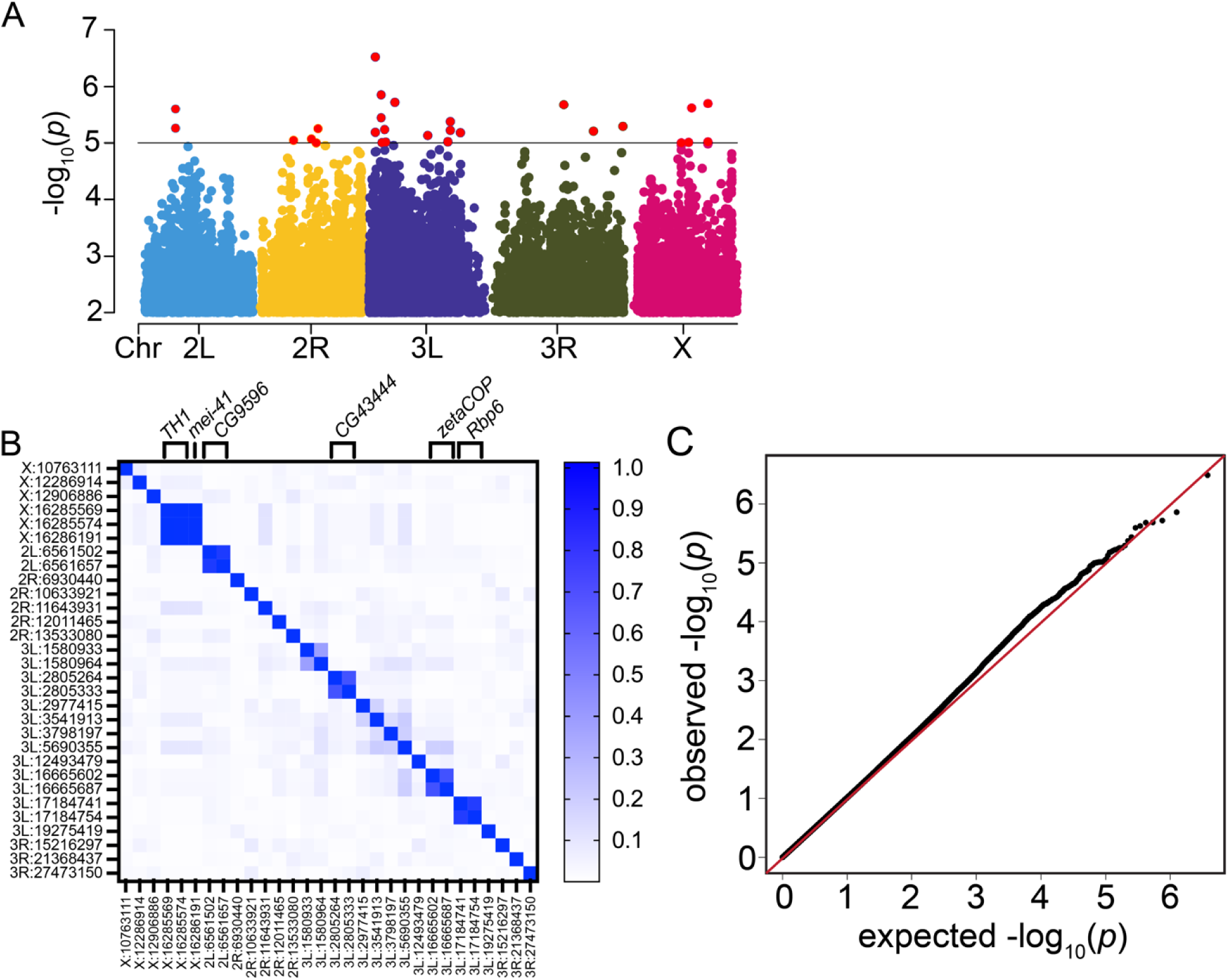
Genome-wide association with paraquat neurodegeneration. (a) –log_10_ P-values for each SNP (MAF ≥ 0.05, < 30% missing) in association with DA neuron viability following paraquat exposure (P-value cutoff of P <0.01). The line indicates SNPs below the *P* <10^-6^ threshold). (B) A pairwise LD heatmap for all 30 associating SNPs (*P* <10^-6^), groups of SNPs in LD are indicated with brackets at the top and annotated by gene, color scale for *r^2^* is shown. (C) A Q-Q plot of –log_10_ *P*-values for the variants tested in the GWAS shows the data fit a normal distribution.

**Table 1.** SNPs associated with DA neuron viability following paraquat exposure.

| Rank | SNP<sup>a</sup> | Candidate gene(s) | P-value | Mixed model<br>P-value | Human ortholog |
| --- | --- | --- | --- | --- | --- |
| 1 | 3L_1580964 | CG13917 | 3.23<sup>-07</sup> | 3.36<sup>-07</sup> | BTBD8 |
| 2 | 3L_2805264 | Tet | 1.38<sup>-06</sup> | 1.36<sup>-06</sup> | TET3 |
| 3 | 3L_5690355 | sif | 1.90<sup>-06</sup> | 1.84<sup>-06</sup> | TIAM1 |
| 4 | X_16285569 | TH1 | 2.05<sup>-06</sup> | 6.85<sup>-06</sup> | NELFCD |
| 5 | 3R_15216297 | CG18493, Ino80 | 2.07<sup>-06</sup> | 2.19<sup>-06</sup> | PRSS16, INO80 |
| 6 | X_12906886 | rad | 2.36<sup>-06</sup> | 1.25<sup>-06</sup> | RAP1GAP2 |
| 7 | 2L_6561657 | Trmt6 | 2.53<sup>-06</sup> | 2.55<sup>-06</sup> | TRMT6 |
| 8 | 3L_2805333 | Tet | 3.63<sup>-06</sup> | 4.45<sup>-06</sup> | TET3 |
| 9 | 3L_17184741 | Rbp6 | 4.22<sup>-06</sup> | 2.50<sup>-06</sup> | MSI2 |
| 10 | 3R_27473150 | kek6 | 5.07<sup>-06</sup> | 1.88<sup>-06</sup> | LRFN1 |
| 11 | 2L_6561502 | Trmt6 | 5.52<sup>-06</sup> | 4.78<sup>-06</sup> | TRMT6 |
| 12 | 2R_12011465 | CG30095 | 5.56<sup>-06</sup> | 4.81<sup>-06</sup> | none |
| 13 | 3L_3541913 | Eip63E | 5.84<sup>-06</sup> | 7.42<sup>-06</sup> | CDK14 |
| 14 | 3L_17184754 | Rbp6 | 5.95<sup>-06</sup> | 3.66<sup>-06</sup> | MSI2 |
| 15 | 3R_21368437 | msi | 6.18<sup>-06</sup> | 1.95<sup>-05</sup> | MSI2 |
| 16 | 3L_1580933 | CG13917 | 6.47<sup>-06</sup> | 6.78<sup>-06</sup> | BTBD8 |
| 17 | 3L_19275419 | Gbs-76A | 6.62<sup>-06</sup> | 6.15<sup>-06</sup> | PPP1R3D |
| 18 | 3L_12493479 | CG10638 | 7.43<sup>-06</sup> | 1.40<sup>-05</sup> | AKR1B15 |
| 19 | 2R_10633921 | Dro | 8.52<sup>-06</sup> | 7.59<sup>-06</sup> | none |
| 20 | 2R_6930440 | luna | 8.96<sup>-06</sup> | 1.67<sup>-05</sup> | KLF7 |
| 21 | 3L_16665687 | zetaCOP | 9.49<sup>-06</sup> | 8.69<sup>-06</sup> | COPZ1 |
| 22 | 3L_16665602 | zetaCOP | 9.61<sup>-06</sup> | 8.91<sup>-06</sup> | COPZ1 |
| 23 | X_16286191 | mei-41 | 9.63<sup>-06</sup> | 1.09<sup>-05</sup> | ATR |
| 24 | X_16285574 | TH1 | 9.63<sup>-06</sup> | 1.09<sup>-05</sup> | NELFCD |
| 25 | 3L_3798197 | CG32264 | 9.67<sup>-06</sup> | 9.36<sup>-06</sup> | PHACTR2 |
| 26 | X_12286914 | intergenic | 9.80<sup>-06</sup> | 1.59<sup>-05</sup> | - |
| 27 | 3L_2977415 | intergenic | 9.92<sup>-06</sup> | 1.13<sup>-05</sup> | - |
| 28 | 2R_11643931 | CG30089 | 1.00<sup>-05</sup> | 9.68<sup>-06</sup> | none |
| 29 | X_10763111 | CG2157 | 1.08<sup>-05</sup> | 9.93<sup>-06</sup> | none |
| 30 | 2R_13533080 | intergenic | 1.12<sup>-05</sup> | 7.42<sup>-06</sup> | - |

SNPs associated at $P < 10^{-6}$ by simple regression or mixed effect model P-values.<sup>a</sup> SNP with most statistically significant association for the candidate gene.

Interestingly, *Phactr2*, a potential mammalian ortholog of *CG32264*, contains a SNP (rs11155313) previously identified as a PD risk factor based on human GWAS and follow-up study in a population-based patient-control series (23). The function of the Phactr family of proteins is not well understood, although they are thought to bind actin and Phactr1 is expressed at high levels in the striatum and enriched at the synapse (24) placing it in a brain region relevant to PD pathophysiology. We used gene ontology (GO) analysis in PANTHER for biological processes overrepresented in paraquat-mediated DA neuron loss associated genes (at *P* <10^-5^) (Table 2). The largest enrichment was for genes with a functional role in synapse organization (5.9 fold enrichment, *P* =7.8×10^-6^, FDR =9.4×10^-3^), including several genes specifically involved in synaptic connection or scaffolding (*caps, prosap, Fife, Nlg4, dpr2* and *dpr15*).

**Table 2.** Biological Process enrichment.

| **GO Biological Process-complete** | **Genome** | **Observed** | **Expected** | **Fold enrichment** | **P-value** | **FDR** |
| --- | --- | --- | --- | --- | --- | --- |
| synapse organization | 191 | 10 | 1.7 | 5.88 | $7.79^{-06}$ | $9.44^{-03}$ |
| cell junction organization | 259 | 11 | 2.3 | 4.77 | $1.96^{-05}$ | $1.42^{-02}$ |
| regulation of multicellular organismal process | 579 | 17 | 5.15 | 3.3 | $1.43^{-05}$ | $1.30^{-02}$ |
| regulation of biological process | 4009 | 60 | 35.68 | 1.68 | $3.38^{-06}$ | $4.91^{-03}$ |
| biological regulation | 4229 | 64 | 37.64 | 1.7 | $9.29^{-07}$ | $3.38^{-03}$ |

Top GO group (sorted by fold enrichment) for Biological Process among the associating genes at $P < 10^{-5}$ . Analysis results are sorted hierarchically for ontology terms within the group. Genome is the number of relevant genes within the *Drosophila* genome ( $n = 13,767$ genes), Expected is the number of genes expected based on the *Drosophila* genome reference list.

### Targeted functional assessment of candidate genes

To identify candidate genes that play a role in resistance to paraquat-mediated DA neurodegeneration, we performed gene-level RNAi studies on all genes nominated at a *P* <10^-6^ SNP association threshold (Table 1) and those annotated with *synapse organization* from our GO enrichment analysis. RNAi lines were unavailable for 7 out of 22 top association genes and for 4 out of 10 synapse organization genes. In total we tested 31 RNAi lines targeting 21 genes (two independent RNAi lines where available), and divided randomly into 3 separate testing cohorts for practical purposes. RNAi lines for each candidate gene were mated to flies harboring the *TH-GAL4* driver to obtain progeny with DA neuron-specific gene knock-down for examining paraquat-induced DA neuron death. As with DGRP testing, we assessed DA neuron viability in flies exposed to 1 mM paraquat continuously for 7d then housed on food without paraquat for an additional 14 days. We observe no significant DA neuron loss for genetic (RNAi vector) control strains exposed to paraquat in each of three experimental cohorts tested, while RNAi for *Gbs-76A*, *TH1*, *luna* and *CG32264* each result in significant DA neuron loss when exposed to paraquat in at least one RNAi line (Fig. 4A), suggesting a phenotypic enhancement and consistent with a role for each gene in contributing towards paraquat resistance. All four genes were nominated from our top associating SNPs (*P* <10^-6^), hence 4 out of 15 nominated genes (27%) tested affect the DA neuron phenotype. The extent of DA neuron loss seen upon knocking-down individual genes is generally modest, suggesting that paraquat-mediated DA neuron loss is likely a polygenic trait not attributable to any single associated gene identified here. In addition to a paraquat-induced phenotype, RNAi to *CG30089* or *fz2* results in significantly reduced numbers of viable DA neurons in the absence of paraquat exposure, raising the possibility that these genes may be required for DA neuron development and/or maintenance across age independent of paraquat.

**Figure 4.**
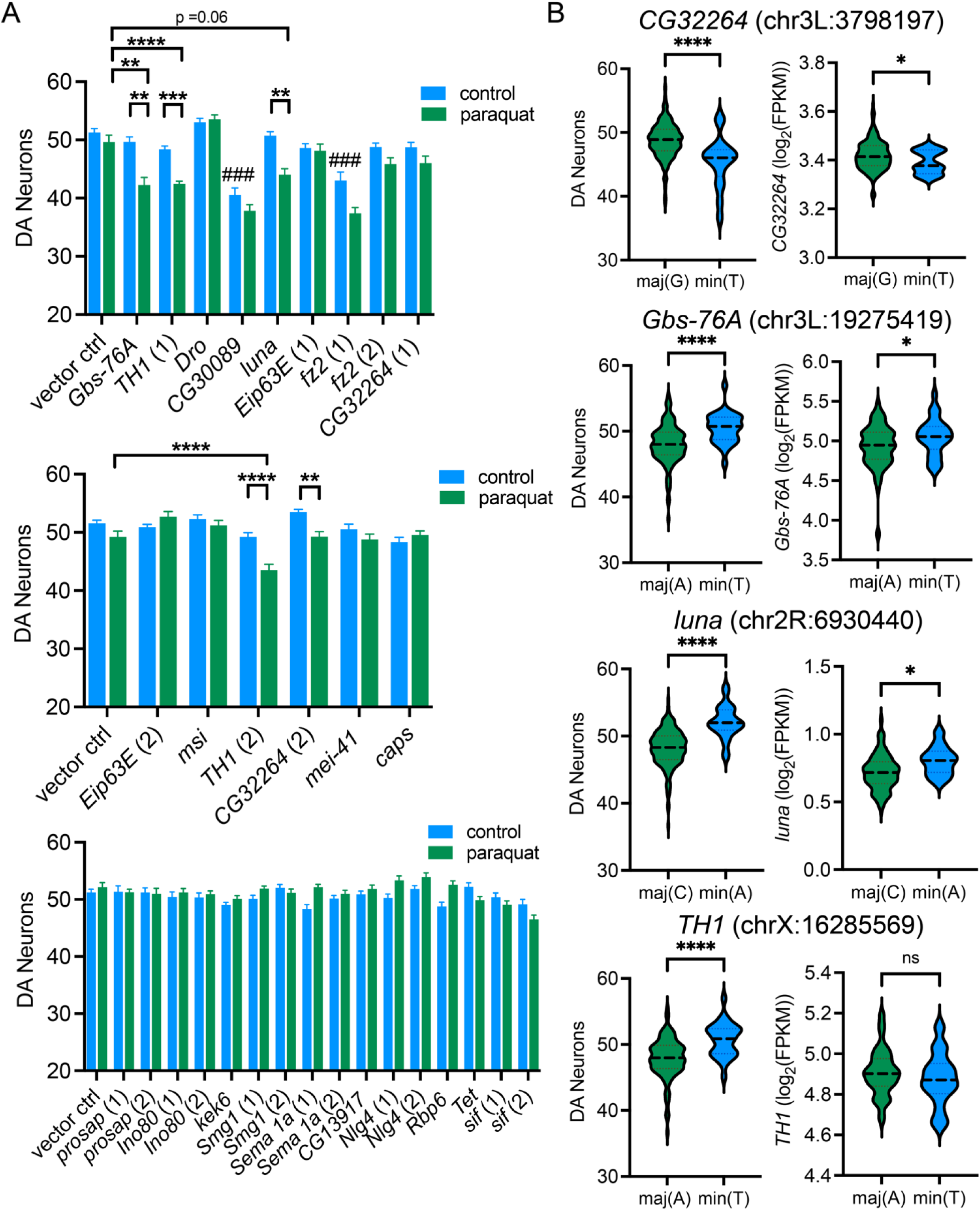
Gene-level testing by RNAi uncovers genes associated with paraquat resistance. (A) 31 RNAi lines collectively targeting knock-down of 21 candidate genes were tested in three experimental groups. DA neuron viability exhibits significant effects of genotype and paraquat exposure and a significant interaction between both factors (individual two-way ANOVAs, P <0.0001(genotype), P <0.01(paraquat), P <0.001(interaction), n=8-15 brains/genotype), Holm-Šídák post-test for effect of RNAi vs genetic control (### P<0.001), effect of paraquat exposure (** P 0.01, *** P <0.001, **** P <0.0001) within line or vs. genetic control. (B) SNP alleles associated with DA neuron viability following paraquat exposure may influence gene expression. DA neuron viability and transcript abundance for candidate genes indicated by RNAi studies are significantly affected by SNP allele for *CG32264*, *Gbs-76A* and *luna* (individual t-tests, * P <0.05, **** P <0.0001). In each case, the SNP allele associated with lower DA neuron viability following paraquat exposure is also associated with lower transcript abundance.

**Figure 5.**
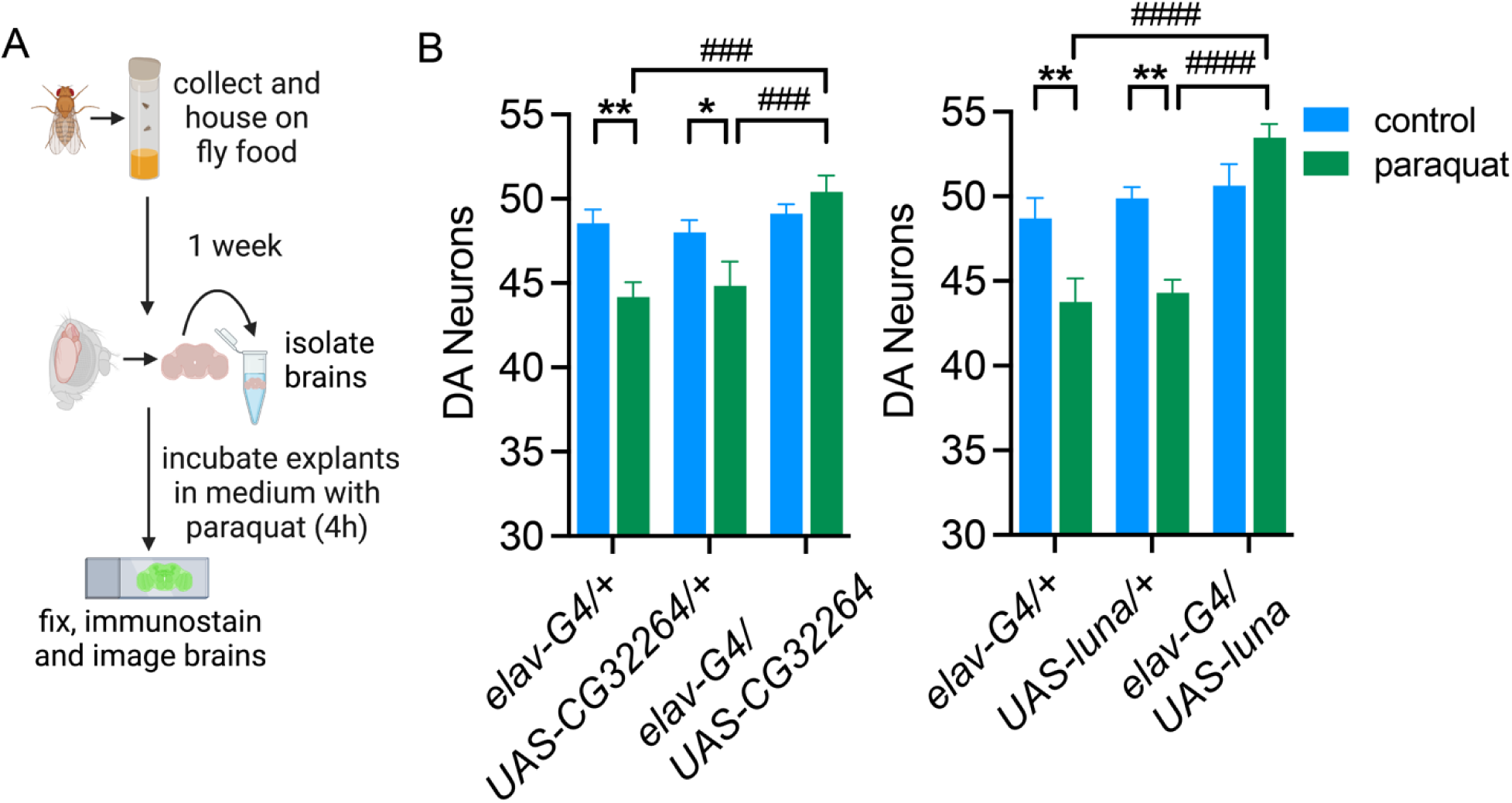
Candidate gene overexpression testing. (A) Scheme illustrating brain explant paraquat exposure. (B) overexpression of *CG32264* or *luna* protects against 100 uM paraquat exposure in brain explants (individual two-way ANOVAs for effect of genotype (P <0.01), paraquat (P <0.05), interaction (P <0.05), n =10-13 brains per genotype, with Tukey’s post-tests for effect of paraquat (* P <0.05, ** P <0.01) and genotype (### P <0.01, #### P <0.0001)).

We used published transcriptomics data for the DGRP to probe for possible effects of associating SNPs in *Gbs-76A*, *TH1*, *luna* and *CG32264* on gene expression and to determine whether any anticipated gene expression changes correlate with results obtained in gene-level RNAi studies. We first parsed paraquat-induced DA neuron death for all tested DGRP strains into minor and major allele harboring strains for the top associating SNP in *Gbs-76A*, *TH1*, *luna* and *CG32264* (Fig. 4B). We next parsed reported DGRP female gene expression data (25) into minor and major allele strains for the same SNPs. This analysis reveals that for *Gbs-76A*, *luna* and *CG32264,* the allele associated with lower DA neuron viability following paraquat exposure is also associated with lower gene expression (Fig. 4B). This links reduced expression with impaired DA neuron integrity following paraquat exposure and agrees with our finding that RNAi-mediated knock-down of each of these genes results in paraquat-induced DA neuron loss. For *TH1*, there is no significant difference in reported transcript levels between minor and major allele carrying strains (Fig. 4B). Hence our limited expression analysis indicates overall robust consistency between the anticipated effect of associating SNPs on gene expression and 3 out of 4 positive RNAi gene candidates.

Since knock-down of *Gbs-76A*, *TH1*, *luna* or *CG32264* results in paraquat-induced DA neuron loss then it might be expected that overexpressing each of these genes would protect DA neurons challenged with paraquat. While our control strains consistently exhibit no significant DA neuron attrition in a paraquat feeding approach, we found that brief incubation of brain explants in culture medium containing paraquat (100 μM) leads to significant DA neuron loss and harnessed this approach to assess putative suppressor effects of candidate gene overexpression. We were able to obtain lines harboring UAS-transgenes for *CG32264* and *luna* while equivalent lines for *Gbs-76A* and *TH1* were unavailable. We crossed these transgenic lines to *elav-GAL4* to drive pan-neuron transgene expression for each candidate gene. Overexpression of *CG32264* or *luna* both prevent paraquat-induced DA neuron loss, while transgenic lines harboring *elav-GAL4* or *UAS-transgenes* alone exhibit neuronal death. Hence, these data support a role for *CG32264* and *luna* in preserving DA neuron survival upon paraquat exposure corroborating evidence from genome-wide association studies based on the anticipated impact of associating polymorphism effects on gene expression and from RNAi studies.

## DISCUSSION

Genetic background frequently influences the expressivity of disease-related phenotypes. Genetics is known to play a major role in PD development, both in monogenic forms of the disease due to mutations in single genes and in idiopathic PD, in which numerous risk variants have been identified via GWAS (9, 26). As environmental factors also play a major role in PD risk, it is plausible that the penetrance of some genetic variants may be modulated by environmental variables through gene-environment interactions. Human studies have revealed several genetic variants that modulate PD risk following pesticide exposures, e.g. *CYP2D67*, *NOS8*, *DAT9*, *Paraoxonase-110* and *ALDH11* (27–31). While some of these genetic insights relate to pesticide detoxification, (e.g. *CYP2D6*), others may begin to provide clues to biological mechanisms, yet, it is difficult to identify alterations in targeted genes that confer pesticide risk in human studies and even more challenging to take a comprehensive, unbiased approach in gene identification. Genetically tractable animal models offer untapped potential for providing new insights into PD etiology through the study of gene-environment interactions.

Here, through a genome-wide analysis and follow-up gene-level studies, we identified several candidate genes providing resistance to DA neurodegeneration caused by exposure to the pesticide paraquat. We used a paraquat concentration that causes a slow-onset attrition of DA neurons without substantial mortality prior to neuronal loss (Fig. 1B, C) and no effect on feeding behavior (Fig. 1D). Since the time elapsed between paraquat exposure and DA neuron assessment in this paradigm involves two weeks of additional aging which is a considerable portion of the average fly lifespan, we considered the possibility that DA neuron phenotypes could be influenced by aging and potentially occur independent of paraquat exposure. However, this scenario is refuted by a low correlation between the number of TH-immunopositive DA neurons observed after paraquat exposure with the number found at the same age in the same DGRP strains tested either in our laboratory or in an independent study (Fig. 2C, D). As DA neurons are uniformly higher in the absence of paraquat exposure, this further argues against variation in DA neuron numbers between DGRP strains being developmental in origin. We found that neurodegeneration is phenotypically divergent over the DGRP collection and is associated with polymorphisms in genes covering a broad range of functions. Our GWAS relies on the association of ∼1.9 million SNPs over 173 DGRP strains, which does not enable sufficient power to identify definitive SNPs associating with paraquat-induced DA neuron loss that survive multiple testing correction. Since this approach increases the chances of identifying false-positive associations, we screened 21 individual genes nominated by these SNPs to encompass a large selection of candidate mediators of paraquat resistance in downstream RNAi studies. Individually silencing four genes, *Gbs-76A*, *TH1*, *luna* and *CG32264* causes significant DA neuron loss upon paraquat exposure consistent with each gene providing a protective effect, albeit subtle, against paraquat neurotoxicity. As noted above, *C32264* is predicted to play a role in actin cytoskeleton organization and is the ortholog of *Phactr2*, previously linked to PD through genetic studies. *Gbs-76A* is predicted to positively regulate glycogen biosynthesis and its putative yeast ortholog, *GIP2*, has been tentatively linked to PD as *GIP2* overexpression was found to suppress alpha synuclein toxicity in yeast (32). TH1 is a constituent of the NELF complex. This is involved in a major checkpoint during transcription called proximal promoter pausing whereby RNA polymerase II pauses downstream of a transcription start site (33). *luna* is a Krueppel-like factor (KLF)-like gene encoding a probable transcription factor (34), which together with *TH1* highlights a potential role for transcriptional regulation in paraquat resistance. While our GO enrichment analysis implicated several genes annotated for a role in synapse organization in paraquat-mediated DA neuron survival, follow-up RNAi assessment did not corroborate this, with the caveat that we were only able to test 6 out of 10 candidate genes in this ontology class due to RNAi line availability. As 0 out of 6 gene candidates tested by RNAi produced a DA neuron phenotype, we currently do not have sufficient evidence in support of synapse organization representing a major functional class of genes relevant to paraquat neurodegeneration. Despite this, we note that the Phactr family of proteins, putatively orthologous to *CG32264* are reportedly enriched at the synapse (24), and as they are thought to interact with the actin cytoskeleton, could conceivably overlap functionally with the sub-class of genes identified from our GWAS that play a role in synaptic connection or scaffolding (*caps, prosap, Fife, Nlg4, dpr2* and *dpr15*). Hence, further investigation into synaptic involvement in paraquat neurotoxicity is warranted from these findings.

Prior comprehensive DGRP gene expression analysis (25) combined with genotype information available for each strain enables a targeted estimation of SNP allele effects on gene expression. For three of four positive candidate genes (*Gbs-76A*, *luna* and *CG32264*), analysis of the top associating SNP indicates that the allele associated with lower DA neuron viability also associates with lower reported gene expression, consistent with our RNAi findings that knocking-down each of these genes results in DA neuron loss. Conversely, *TH1* does not exhibit allele specific expression for the top SNP associated with paraquat neurodegeneration. Notably, the gene expression profiling used for this analysis was conducted on 2-day-old flies (25) while our DA neuron phenotype was measured in 21-day-old flies. Hence, if *TH1* exhibits age-related expression changes, it’s possible that these may interact with the associating SNP and if so, this would not be identified in our analysis. We did not perform a comprehensive assessment of expression quantitative trait loci (eQTLs) across the fly genome here as this was beyond the scope of our study. Instead, we focused on a limited assessment targeting individual genes implicated by RNAi studies and harboring an associating SNP of interest in order to better characterize consistency between our GWAS and RNAi findings.

To determine whether increased expression of candidate paraquat resistance genes can protect against paraquat toxicity, we examined DA neuron loss in brain explants treated with paraquat. The use of brain explants allowed us to circumvent the lack of paraquat toxicity seen in outcrossed control strains (in contrast to a subset of inbred DGRP strains). We were able to obtain transgenic lines under UAS regulation for *luna* and *CG32264*, and overexpression of either gene prevents paraquat-induced neurodegeneration, underscoring their ability to promote paraquat resistance.

Our study builds upon the demonstrated utility of the DGRP to yield new insight into the genetics of disease-associated traits. Prior studies have revealed genes associated with neurodegeneration related to an Rh1 mutation in retinitis pigmentosa (35), Amyloid β42 and tau toxicity in Alzheimer’s disease (16), and mutant LRRK2 expression in Parkinson’s disease (17). While the DGRP have been used to study the genetic basis of resistance to Parkinson’s disease-linked pesticides such as paraquat (21) and DDT (36), to our knowledge this is the first study focused on a neurodegenerative phenotype caused by any PD-linked pesticide. Given their genetic tractability and ease of controlled pesticide exposure, we believe that *Drosophila* can provide a powerful gene discovery platform for studying gene-environment interactions relevant to PD. Moreover, the broad neurodegenerative phenotype divergence in the DGRP found in this study and prior studies can be leveraged to gain mechanistic insight into exposure to pesticides or other environmental toxicants that do not produce robust phenotypes in standard animal control strains.

## ACKNOWLEDGMENTS

We thank the Oregon Health and Science University Advanced Light Microscopy Core (RRID:SCR_009961) for expert technical assistance. This work was supported by Department of Defense Grant PD200015 to I.M.

## AUTHOR CONTRIBUTIONS

S.V-C. – Conceptualization, Investigation, Formal analysis, Writing manuscript A.A-B. - Investigation, Formal analysis, Writing manuscript I.M. - Conceptualization, Investigation, Formal analysis, Writing manuscript, Supervision, Funding acquisition.

## MATERIALS AND METHODS

### Drosophila stocks and culture

173 DGRP lines used in the study (Table S1) were obtained from the Bloomington *Drosophila* Stock Center. The following lines carrying GAL4, RNAi or cDNA constructs under UAS enhancer control were obtained from the Bloomington *Drosophila* stock center (line numbers in parenthesis). GAL4 lines: *TH-GAL4* (8848), *elav^C155^-GAL4* (458); RNAi lines: *CG13917* (42903), *Tet* (62280), *sif* (25789, 61934), *kek6* (61212), *ino80* (33708, 37473), *Rbp6* (61324), *Gbs-76A* (42551), *TH1* (38934, 42931), *Eip63E* (28901, 34075), *Dro* (67223), *CG32264* (28340, 64966), *CG30089* (28321), *mei-41* (41934), *luna* (27084), *msi* (55152) and the VALIUM 10/20 vector control *UAS-LUC.VALIUM10* (35788); transgenic overexpression lines: *luna* (56814). *UAS-CG32264* (F000497) was obtained from FlyORF. All flies were reared and aged at 25°C/60% relative humidity under a 12 h light-dark cycle on standard cornmeal medium.

### Paraquat exposure - feeding

Paraquat (or water control) was vigorously mixed into molten standard fly food to a 1 mM final concentration and the food was immediately cooled and stored at 4°C until use (<1 week). At this paraquat dose, we see no effect on feeding behavior (Fig. 1D). Female adult flies (average 2 days-old) were collected and transferred to vials containing paraquat-supplemented food or control vials pre-warmed to RT. Flies were housed on paraquat-supplemented food for 7 days, with transfer to fresh paraquat food at day 4, then subsequently transferred to standard fly food without paraquat for an additional 14 days. Flies were processed for dopamine neuron immunostaining as described below.

### Paraquat exposure – brain explants

Adult female brains were isolated and transferred to Schneider’s *Drosophila* medium (formulated for *Drosophila* cells and tissues and supplemented with L-glutamine and sodium bicarbonate), using reported methods (37). Schneider’s medium was then replaced with 100 μM paraquat in Schneider’s medium and the brains were incubated on a thermomixer (25 °C, 300 rpm) for 4 h, washed in 1 ml of Schneider’s medium, fixed and permeabilized in 4% PFA in 0.3% PBS-T for 20 min at RT and then processed for DA neuron immunostaining (see below) from the blocking step onwards.

### Dopamine neuron immunostaining

Brains (10-20/genotype) were harvested from heads pre-fixed in 4% PFA in 0.3% PBS-T for 15 min. Brains were fixed, and permeabilized with 4% PFA in 0.3% PBS-T, pH 7.4 for 20 min at RT, then blocked in 2.5% normal donkey serum for 1 h at room temperature. Brains were incubated in anti-tyrosine hydroxylase (1:1000, Immunostar) for three nights on a nutator at 4°C. After extensive washing, brains were then incubated with Alexa Fluor 488 secondary antibody (1:2000) for 3 nights at 4°C. Brains were washed extensively and whole-mounted in SlowFade Gold Antifade mounting medium. Confocal z-stack images of the stained brains were acquired on Zeiss LSM 900 at a 1 μm slice interval and dopamine neurons with any detectable tyrosine hydroxylase were counted. Dopamine neurons belonging to the protocerebral posterior lateral 1 (PPL1), protocerebral posterior medial 1/2 (PPM1/2), and protocerebral posterior medial 3 (PPM3) clusters were counted.

### Genome-wide association

DA neuron viability in 173 DGRP lines was tested in 12 blocks, with each block including an established paraquat-sensitive line (RAL-911) and a paraquat-resistant line (RAL-821) as internal controls. We estimated broad-sense heritability within each of the 12 blocks that contained 14-20 randomly picked DGRP lines. For each experimental block, an independent estimate of heritability was derived in an ANOVA by *σ_L_*^2^ / (*σ_L_*^2^ + *σ_E_*^2^), where *σ_L_*^2^ represents the between-line variance of (non-transformed) DA neuron counts and *σ_E_*^2^ is the within-line variance. Overall mean heritability was calculated across all twelve individual block estimates.

Raw DA neuron counts (for PPM1/2, PPM3 and PPL1 clusters combined) displayed variance across blocks (Mixed Model (random effects), block variance = 3.47 (32%), Residual variance = 7.29 (68%)). Block effects were corrected using residuals from internal control genotypes (RAL-911 and RAL-821 combined mean) centered to mean = 0. Block-corrected phenotype data were uploaded to the DGRP portal (http://dgrp.gnets.ncsu.edu/) which uses PLINK, FaST-LMM, SnpEff and R to perform genome-wide association and generate output files. Variants were filtered for minor allele frequency (≥ 0.05), missingness (<30%) and non-biallelic sites were removed. A total of 1,894,786 SNPs were included in the analysis and DA neuron counts for 173 DGRP lines were regressed on each SNP. Quantile-quantile (Q-Q) plot analysis indicated normally distributed data (Fig. 3C). Regression analysis revealed that neither *Wolbachia* infection status nor five major chromosomal inversions found in the DGRP were significant covariates (Table S3). Pairwise linkage disequilibrium (LD) among top associated SNPs (*P* <10^-6^) suggested that SNPs in LD (*r^2^* ≥0.5) were within the same gene boundaries in all but one case consistent with overall low LD. SNPs chrX:16285574 in *TH1* and chrX:16286191 in *mei-41* are in LD (*r^2^* =1) but separated by only ∼600 bp. SNPs associating at p <0.01 were Genes were identified from SNP coordinates using the BDGP R54/dm3 genome build. A SNP was assigned to a gene if it was within the gene’s transcription boundaries or no more than 1,000 bp upstream/downstream of those boundaries. Gene IDs for SNPs at *P* <10^-5^ were analyzed in the PANTHER 19.0 database for GO biological process-complete enrichment against all *Drosophila* genes in the database using Fisher’s exact test.

### Statistical analysis

Unless otherwise described in the methods, data were analyzed by Student’s t-test, ANOVA or two-way ANOVA with correction for multiple comparisons.

### Data availability

SNP location data in the DGRP collection (Freeze 2.0 calls) can be downloaded from the DGRP data portal (http://dgrp2.gnets.ncsu.edu/data/website/dgrp2.tgeno) and are additionally deposited on the GSA Figshare portal along with DGRP genotype data and raw phenotype association data.

## Notes

### Competing Interest Statement

The authors have declared no competing interest.

