## Supplemental Tables S1-S3 for "Genome-wide analysis reveals genes mediating resistance to paraquat neurodegeneration in *Drosophila*"

**Table S1.** DGRP lines, dopamine neuron counts, strain *Wolbachia* infection status and survivorship at end of 21-day paraquat exposure plus aging period.

| **DGRP line**  **number** | **DA neurons (mean)** | ***Wolbachia* infection status** | **Survival at 21 days (%)** |
| --- | --- | --- | --- |
| 21 | 48.9 | Y | 100 |
| 26 | 49.3 | N | 96 |
| 28 | 41.4 | N | 100 |
| 31 | 52.4 | N | 100 |
| 32 | 50.5 | N | 100 |
| 40 | 51.0 | Y | 100 |
| 41 | 51.1 | N | 100 |
| 42 | 54.1 | N | 100 |
| 45 | 50.2 | N | 100 |
| 48 | 52.7 | Y | 100 |
| 57 | 51.0 | N | 98 |
| 59 | 52.5 | N | 100 |
| 69 | 51.9 | Y | 96 |
| 73 | 50.7 | Y | 96 |
| 75 | 51.9 | Y | 100 |
| 83 | 46.3 | N | 98 |
| 85 | 54.5 | N | 100 |
| 88 | 47.4 | N | 100 |
| 91 | 46.2 | N | 100 |
| 93 | 50.9 | N | 100 |
| 100 | 49.1 | Y | 100 |
| 101 | 47.9 | N | 100 |
| 105 | 49.3 | N | 100 |
| 109 | 47.6 | N | 100 |
| 129 | 49.4 | N | 100 |
| 136 | 53.2 | Y | 100 |
| 142 | 50.5 | Y | 100 |
| 149 | 48.3 | Y | 100 |
| 153 | 49.4 | Y | 96 |
| 161 | 48.5 | N | 98 |
| 177 | 47.8 | N | 94 |
| **DGRP line** | **DA neurons (mean)** | ***Wolbachia* infection status** | **Survival at 21 days (%)** |
| 181 | 49.2 | Y | 98 |
| 189 | 45.9 | Y | 94 |
| 195 | 47.0 | N | 92 |
| 208 | 49.9 | N | 100 |
| 217 | 49.4 | N | 100 |
| 227 | 44.1 | Y | 82 |
| 228 | 42.8 | N | 96 |
| 229 | 39.9 | N | 92 |
| 235 | 48.8 | N | 98 |
| 239 | 57.0 | N | 90 |
| 256 | 46.4 | Y | 98 |
| 280 | 44.6 | Y | 100 |
| 287 | 48.8 | Y | 98 |
| 301 | 45.9 | N | 98 |
| 303 | 47.7 | N | 98 |
| 304 | 46.9 | Y | 98 |
| 306 | 44.5 | Y | 100 |
| 309 | 46.1 | N | 98 |
| 310 | 50.8 | Y | 96 |
| 313 | 51.0 | N | 98 |
| 315 | 47.1 | N | 96 |
| 317 | 49.2 | Y | 98 |
| 318 | 46.4 | Y | 62 |
| 319 | 47.8 | Y | 100 |
| 320 | 46.9 | Y | 100 |
| 321 | 49.2 | Y | 100 |
| 324 | 48.8 | N | 96 |
| 332 | 47.1 | N | 92 |
| 335 | 50.1 | Y | 100 |
| 336 | 51.6 | Y | 90 |
| 338 | 46.1 | Y | 96 |
| 340 | 45.6 | Y | 100 |
| 348 | 47.4 | N | 98 |
| 350 | 50.7 | N | 100 |
| 355 | 49.7 | Y | 100 |
| 357 | 46.2 | N | 100 |
| **DGRP line** | **DA neurons (mean)** | ***Wolbachia* infection status** | **Survival at 21 days (%)** |
| 358 | 50.0 | N | 100 |
| 359 | 50.1 | N | 100 |
| 360 | 50.7 | Y | 100 |
| 361 | 46.4 | Y | 98 |
| 362 | 48.1 | Y | 100 |
| 370 | 49.4 | Y | 98 |
| 371 | 50.8 | N | 100 |
| 373 | 48.3 | N | 100 |
| 374 | 47.7 | Y | 98 |
| 375 | 47.2 | N | 100 |
| 377 | 47.6 | N | 96 |
| 379 | 49.3 | N | 100 |
| 380 | 48.6 | Y | 96 |
| 381 | 54.2 | N | 96 |
| 383 | 49.4 | Y | 98 |
| 385 | 49.1 | N | 80 |
| 386 | 48.5 | N | 92 |
| 390 | 50.1 | N | 84 |
| 391 | 49.3 | N | 100 |
| 392 | 47.8 | N | 96 |
| 397 | 50.8 | Y | 100 |
| 399 | 36.7 | N | 100 |
| 405 | 46.5 | Y | 98 |
| 406 | 47.7 | N | 96 |
| 426 | 52.0 | N | 100 |
| 427 | 50.9 | N | 100 |
| 439 | 52.2 | N | 98 |
| 440 | 51.9 | Y | 100 |
| 441 | 55.4 | Y | 100 |
| 443 | 52.5 | N | 96 |
| 461 | 49.9 | Y | 88 |
| 486 | 53.7 | Y | 98 |
| 491 | 50.7 | N | 100 |
| 492 | 52.1 | N | 100 |
| 502 | 46.1 | N | 100 |
| 505 | 51.5 | Y | 98 |
| **DGRP line** | **DA neurons (mean)** | ***Wolbachia* infection status** | **Survival at 21 days (%)** |
| 508 | 49.6 | N | 90 |
| 509 | 48.5 | N | 98 |
| 513 | 52.4 | Y | 100 |
| 517 | 52.0 | N | 100 |
| 528 | 51.4 | Y | 92 |
| 530 | 44.6 | Y | 100 |
| 531 | 49.9 | Y | 100 |
| 535 | 46.0 | Y | 96 |
| 551 | 49.9 | Y | 98 |
| 555 | 45.9 | Y | 94 |
| 559 | 48.1 | N | 100 |
| 563 | 48.0 | N | 98 |
| 566 | 50.5 | N | 98 |
| 584 | 49.7 | Y | 100 |
| 589 | 52.5 | Y | 90 |
| 595 | 46.5 | Y | 98 |
| 596 | 47.3 | N | 98 |
| 627 | 47.1 | N | 100 |
| 630 | 48.1 | N | 100 |
| 634 | 48.3 | Y | 100 |
| 639 | 51.1 | Y | 94 |
| 646 | 51.3 | Y | 98 |
| 703 | 46.7 | N | 100 |
| 705 | 45.1 | Y | 100 |
| 707 | 49.9 | Y | 98 |
| 712 | 46.8 | Y | 100 |
| 714 | 47.0 | N | 100 |
| 716 | 45.3 | Y | 100 |
| 721 | 52.4 | Y | 100 |
| 730 | 49.3 | Y | 92 |
| 732 | 48.0 | N | 98 |
| 737 | 49.3 | Y | 100 |
| 738 | 47.9 | Y | 100 |
| 748 | 41.5 | Y | 100 |
| 757 | 50.4 | N | 74 |
| 765 | 49.1 | N | 90 |
| **DGRP line** | **DA neurons (mean)** | ***Wolbachia* infection status** | **Survival at 21 days (%)** |
| 774 | 49.9 | N | 94 |
| 776 | 48.0 | Y | 100 |
| 786 | 39.2 | Y | 98 |
| 787 | 49.1 | Y | 92 |
| 790 | 46.5 | Y | 98 |
| 796 | 47.4 | Y | 100 |
| 799 | 45.2 | N | 98 |
| 801 | 44.0 | Y | 100 |
| 802 | 45.7 | Y | 96 |
| 804 | 48.0 | Y | 96 |
| 808 | 50.5 | N | 94 |
| 810 | 51.9 | N | 98 |
| 812 | 49.1 | N | 98 |
| 818 | 46.5 | Y | 100 |
| 819 | 45.7 | Y | 88 |
| 820 | 45.3 | Y | 100 |
| 821 | 49.0 | Y | 100 |
| 822 | 47.4 | Y | 100 |
| 832 | 47.9 | Y | 100 |
| 837 | 42.8 | Y | 100 |
| 843 | 45.1 | N | 96 |
| 850 | 46.0 | Y | 100 |
| 852 | 48.3 | Y | 100 |
| 853 | 39.1 | Y | 100 |
| 855 | 42.3 | Y | 100 |
| 857 | 47.4 | N | 96 |
| 859 | 46.5 | Y | 100 |
| 861 | 48.7 | Y | 96 |
| 879 | 47.7 | Y | 100 |
| 882 | 47.5 | Y | 100 |
| 884 | 48.6 | Y | 100 |
| 890 | 50.2 | Y | 100 |
| 900 | 47.2 | N | 100 |
| 911 | 42.7 | N | 80 |

**Table S2.** Extended list of associating SNP (*P* <10^-5^). MAF, minor allele frequency. Gene(s) is the gene boundaries ± 1 kb where each SNP is located.

| **SNP ID** | **MAF** | **Minor Allele Count** | **Major Allele Count** | ***P*-value** | **Mixed model**  ***P*-value** | **Gene(s)** |
| --- | --- | --- | --- | --- | --- | --- |
| 3L_1580964_SNP | 0.08671 | 15 | 158 | 3.23E-07 | 3.36E-07 | *CG13917* |
| X_12906886_SNP | 0.08187 | 14 | 157 | 2.36E-06 | 1.25E-06 | *rad* |
| 3L_2805264_SNP | 0.2706 | 46 | 124 | 1.38E-06 | 1.36E-06 | *Tet* |
| 3L_5690355_SNP | 0.05814 | 10 | 162 | 1.91E-06 | 1.84E-06 | *sif* |
| 3R_27473150_SNP | 0.3101 | 49 | 109 | 5.07E-06 | 1.88E-06 | *kek6* |
| 3R_15216297_SNP | 0.08497 | 13 | 140 | 2.07E-06 | 2.19E-06 | *Ino80, CG18493* |
| 3L_17184741_SNP | 0.3774 | 60 | 99 | 4.22E-06 | 2.50E-06 | *Rbp6* |
| 2L_6561657_SNP | 0.07059 | 12 | 158 | 2.53E-06 | 2.55E-06 | *Trmt6* |
| 3L_17184754_SNP | 0.375 | 60 | 100 | 5.95E-06 | 3.66E-06 | *Rbp6* |
| 3L_2805333_SNP | 0.2761 | 45 | 118 | 3.63E-06 | 4.45E-06 | *Tet* |
| 2L_6561502_SNP | 0.06587 | 11 | 156 | 5.52E-06 | 4.78E-06 | *Trmt6* |
| 2R_12011465_SNP | 0.1779 | 29 | 134 | 5.56E-06 | 4.81E-06 | *CG30095* |
| 3L_19275419_SNP | 0.1918 | 28 | 118 | 6.62E-06 | 6.15E-06 | *Gbs-76A* |
| 3L_1580933_SNP | 0.06433 | 11 | 160 | 6.47E-06 | 6.78E-06 | *CG13917* |
| X_16285569_SNP | 0.06433 | 11 | 160 | 2.05E-06 | 6.85E-06 | *TH1* |
| 3L_3541913_SNP | 0.07738 | 13 | 155 | 5.84E-06 | 7.42E-06 | *Eip63E* |
| 2R_10633921_SNP | 0.06509 | 11 | 158 | 8.52E-06 | 7.59E-06 | *Dro* |
| 3L_16665687_SNP | 0.1353 | 23 | 147 | 9.49E-06 | 8.69E-06 | *zetaCOP* |
| 3L_16665602_SNP | 0.1446 | 24 | 142 | 9.61E-06 | 8.91E-06 | *zetaCOP* |
| 3L_3798197_SNP | 0.1053 | 18 | 153 | 9.67E-06 | 9.36E-06 | *CG32264* |
| 2R_11643931_SNP | 0.05233 | 9 | 163 | 1.00E-05 | 9.68E-06 | *CG30089* |
| X_10763111_SNP | 0.4269 | 73 | 98 | 1.09E-05 | 9.93E-06 | *CG2157* |
| X_10763113_SNP | 0.4176 | 71 | 99 | 1.00E-05 | 1.03E-05 | *CG2157* |
| 3L_5417456_SNP | 0.3012 | 50 | 116 | 1.11E-05 | 1.08E-05 | *dip-delta* |
| X_16285574_SNP | 0.06358 | 11 | 162 | 9.63E-06 | 1.09E-05 | *TH1* |
| X_16286191_SNP | 0.06358 | 11 | 162 | 9.63E-06 | 1.09E-05 | *mei-41* |
| 3L_1580822_SNP | 0.07602 | 13 | 158 | 1.44E-05 | 1.13E-05 | *CG13917* |
| X_10763015_SNP | 0.432 | 73 | 96 | 1.33E-05 | 1.13E-05 | *CG2157* |
| 2L_9153038_SNP | 0.05848 | 10 | 161 | 1.16E-05 | 1.18E-05 | *CG32982* |
| X_16285617_SNP | 0.06433 | 11 | 160 | 1.04E-05 | 1.30E-05 | *mei-42* |
| 3R_27187629_SNP | 0.05357 | 9 | 159 | 1.49E-05 | 1.36E-05 | *CG34347* |
| 3L_12493479_SNP | 0.1369 | 23 | 145 | 7.43E-06 | 1.40E-05 | *CG10638* |
| 3L_1752424_SNP | 0.3393 | 57 | 111 | 1.42E-05 | 1.50E-05 | *Cht2* |
| 2R_6930440_SNP | 0.07602 | 13 | 158 | 8.96E-06 | 1.67E-05 | *luna* |
| 3L_16665685_SNP | 0.1404 | 24 | 147 | 1.72E-05 | 1.67E-05 | *zetaCOP* |
| 2R_20297890_SNP | 0.4658 | 75 | 86 | 1.40E-05 | 1.69E-05 | *SerT* |
| **SNP ID** | **MAF** | **Minor Allele Count** | **Major Allele Count** | ***P*-value** | **Mixed model**  ***P*-value** | **Gene(s)** |
| X_10763007_SNP | 0.4438 | 75 | 94 | 2.15E-05 | 1.80E-05 | *CG2157* |
| 2R_6290647_SNP | 0.1928 | 32 | 134 | 2.26E-05 | 1.84E-05 | *SLO2* |
| 3L_874035_SNP | 0.05357 | 9 | 159 | 4.41E-05 | 1.88E-05 | *Usp10* |
| X_21216991_SNP | 0.1529 | 26 | 144 | 1.95E-05 | 1.92E-05 | *dod* |
| 3R_21368437_SNP | 0.122 | 20 | 144 | 6.18E-06 | 1.95E-05 | *msi* |
| X_21217103_SNP | 0.152 | 26 | 145 | 1.56E-05 | 1.96E-05 | *dod* |
| 3R_7121887_SNP | 0.2436 | 38 | 118 | 1.48E-05 | 1.97E-05 | *CG31386* |
| 3R_7552952_SNP | 0.129 | 20 | 135 | 7.13E-05 | 2.00E-05 | *CG12594* |
| 3R_19690443_SNP | 0.06452 | 10 | 145 | 1.79E-05 | 2.02E-05 | *Pli* |
| 3L_2645560_SNP | 0.1647 | 28 | 142 | 2.14E-05 | 2.03E-05 | *CG1143* |
| 3L_5687448_SNP | 0.07738 | 13 | 155 | 2.55E-05 | 2.11E-05 | *sif* |
| 3L_14920208_SNP | 0.2249 | 38 | 131 | 1.32E-05 | 2.14E-05 | *CG42247* |
| X_10763021_SNP | 0.4345 | 73 | 95 | 2.59E-05 | 2.14E-05 | *CG2157* |
| 3L_1752415_SNP | 0.3252 | 53 | 110 | 2.90E-05 | 2.21E-05 | *Cht2* |
| 2R_9959832_SNP | 0.454 | 74 | 89 | 2.82E-05 | 2.29E-05 | *Prosap* |
| X_10763023_SNP | 0.4379 | 74 | 95 | 2.69E-05 | 2.34E-05 | *CG2157* |
| 2R_20998268_SNP | 0.1479 | 25 | 144 | 3.24E-05 | 2.34E-05 | *Tkr* |
| 3L_2686416_SNP | 0.06433 | 11 | 160 | 2.24E-05 | 2.40E-05 | *fife* |
| 3L_3735797_SNP | 0.3452 | 58 | 110 | 2.24E-05 | 2.45E-05 | *CG32264* |
| 2R_20765144_SNP | 0.06509 | 11 | 158 | 2.65E-05 | 2.49E-05 | *Atf-2* |
| 3L_2686418_SNP | 0.06433 | 11 | 160 | 2.26E-05 | 2.49E-05 | *fife* |
| 2R_12113824_SNP | 0.36 | 54 | 96 | 3.14E-05 | 2.51E-05 | *CG15706* |
| 3L_5707466_SNP | 0.1257 | 21 | 146 | 2.72E-05 | 2.53E-05 | *sif* |
| 3L_19318809_SNP | 0.05952 | 10 | 158 | 2.40E-05 | 2.56E-05 | *pip* |
| 3L_6555528_SNP | 0.1012 | 17 | 151 | 2.50E-05 | 2.59E-05 | *MCU* |
| X_12612548_SNP | 0.05263 | 9 | 162 | 2.55E-05 | 2.61E-05 | *Smr* |
| 3L_4258106_SNP | 0.2349 | 39 | 127 | 2.38E-05 | 2.64E-05 | *CG15011* |
| 3R_25726284_SNP | 0.2195 | 36 | 128 | 3.07E-05 | 2.65E-05 | *CG31038* |
| 2L_10557149_SNP | 0.08824 | 15 | 155 | 2.09E-05 | 2.67E-05 | *Trim9* |
| 3L_3735803_SNP | 0.3529 | 60 | 110 | 2.58E-05 | 2.85E-05 | *CG32264* |
| 2L_10853040_SNP | 0.4774 | 74 | 81 | 6.36E-05 | 2.92E-05 | *CG17086* |
| X_10762966_SNP | 0.381 | 64 | 104 | 3.48E-05 | 2.92E-05 | *CG2157* |
| 3L_4093931_SNP | 0.2353 | 40 | 130 | 2.68E-05 | 3.14E-05 | *CG14990* |
| 3L_3542634_SNP | 0.0875 | 14 | 146 | 4.33E-05 | 3.22E-05 | *Eip63E* |
| 3L_17187431_SNP | 0.2778 | 45 | 117 | 3.70E-05 | 3.36E-05 | *CG32169* |
| 2L_9336482_SNP | 0.08824 | 15 | 155 | 3.07E-05 | 3.46E-05 | *Eaat1* |
| X_21217294_SNP | 0.1579 | 27 | 144 | 3.26E-05 | 3.49E-05 | *dod* |
| 3R_15217975_SNP | 0.1284 | 19 | 129 | 6.26E-05 | 3.52E-05 | *Ino80* |
| 3L_14086940_SNP | 0.3937 | 63 | 97 | 5.09E-05 | 3.55E-05 | *Fbp1* |
| **SNP ID** | **MAF** | **Minor Allele Count** | **Major Allele Count** | ***P*-value** | **Mixed model**  ***P*-value** | **Gene(s)** |
| 3L_1580820_SNP | 0.07018 | 12 | 159 | 3.70E-05 | 3.59E-05 | *CG13917* |
| 3L_4056192_SNP | 0.1538 | 26 | 143 | 3.82E-05 | 3.69E-05 | *CG1136* |
| 3L_5428668_SNP | 0.2083 | 35 | 133 | 3.81E-05 | 3.70E-05 | *CG34391* |
| 3L_3735862_SNP | 0.3393 | 57 | 111 | 3.11E-05 | 3.70E-05 | *CG32264* |
| 2R_11640995_SNP | 0.06471 | 11 | 159 | 3.88E-05 | 3.79E-05 | *CG30089* |
| 3L_4874844_SNP | 0.09942 | 17 | 154 | 3.67E-05 | 3.79E-05 | *CG17150* |
| 3L_5688790_SNP | 0.1916 | 32 | 135 | 3.76E-05 | 3.82E-05 | *sif* |
| 2L_10544571_SNP | 0.1056 | 17 | 144 | 4.22E-05 | 3.84E-05 | *Trim9* |
| X_6728261_SNP | 0.1686 | 29 | 143 | 6.38E-05 | 3.91E-05 | *Smg1* |
| 3L_5570229_SNP | 0.05233 | 9 | 163 | 3.98E-05 | 3.91E-05 | *Msr-110* |
| X_21217309_SNP | 0.1538 | 26 | 143 | 3.86E-05 | 3.95E-05 | *dod* |
| 3L_12242152_SNP | 0.256 | 43 | 125 | 4.32E-05 | 4.01E-05 | *CG42318* |
| 2R_17689260_SNP | 0.1369 | 23 | 145 | 4.30E-05 | 4.03E-05 | *tpr* |
| 3R_15708672_SNP | 0.2152 | 34 | 124 | 5.65E-05 | 4.05E-05 | *Pk92B* |
| 2L_10853059_SNP | 0.1012 | 17 | 151 | 3.95E-05 | 4.07E-05 | *CG17086* |
| 3L_13882330_SNP | 0.05128 | 8 | 148 | 8.01E-05 | 4.08E-05 | *CG34400* |
| 2R_9974586_SNP | 0.06548 | 11 | 157 | 4.45E-05 | 4.12E-05 | *Prosap* |
| 3L_5626591_SNP | 0.07692 | 13 | 156 | 3.94E-05 | 4.22E-05 | *Blimp-1* |
| X_4321990_SNP | 0.09677 | 15 | 140 | 4.39E-05 | 4.22E-05 | *bi* |
| 2L_8049843_SNP | 0.425 | 68 | 92 | 4.57E-05 | 4.23E-05 | *Megf8* |
| 2R_16367455_SNP | 0.06471 | 11 | 159 | 3.53E-05 | 4.23E-05 | *side-VIII* |
| 2R_16367451_SNP | 0.06471 | 11 | 159 | 4.11E-05 | 4.32E-05 | *side-VIII* |
| 2R_20398680_SNP | 0.05294 | 9 | 161 | 3.15E-05 | 4.33E-05 | *Letm1* |
| 2L_8049765_SNP | 0.4839 | 75 | 80 | 2.83E-05 | 4.45E-05 | *Megf8* |
| 3L_15938867_SNP | 0.08929 | 15 | 153 | 5.07E-05 | 4.47E-05 | *Pka-C3* |
| 2R_9974612_SNP | 0.06587 | 11 | 156 | 5.09E-05 | 4.48E-05 | *Prosap* |
| 3L_6026172_SNP | 0.05917 | 10 | 159 | 4.71E-05 | 4.49E-05 | *CG10477* |
| 3L_3566304_SNP | 0.1686 | 29 | 143 | 4.18E-05 | 4.52E-05 | *Eip63E* |
| 2L_10963729_SNP | 0.189 | 31 | 133 | 5.54E-05 | 4.57E-05 | *dpr2* |
| 2R_10634726_SNP | 0.142 | 24 | 145 | 3.96E-05 | 4.63E-05 | *AttA* |
| 3L_16149182_SNP | 0.07186 | 12 | 155 | 4.97E-05 | 4.66E-05 | *CG5151* |
| 2R_6290627_SNP | 0.1951 | 32 | 132 | 6.02E-05 | 4.67E-05 | *SLO2* |
| X_12612245_SNP | 0.07097 | 11 | 144 | 6.99E-05 | 4.69E-05 | *Smr* |
| 3R_7289519_SNP | 0.05233 | 9 | 163 | 4.52E-05 | 4.78E-05 | *CG17230* |
| 3L_6026187_SNP | 0.05952 | 10 | 158 | 5.28E-05 | 4.79E-05 | *CG10477* |
| X_21228531_SNP | 0.1279 | 22 | 150 | 4.46E-05 | 4.81E-05 | *sol* |
| 2R_18315219_SNP | 0.3787 | 64 | 105 | 4.75E-05 | 4.86E-05 | *ppk12* |
| X_20944680_SNP | 0.4793 | 81 | 88 | 5.13E-05 | 4.92E-05 | *bves* |
| 3L_4515851_SNP | 0.4529 | 77 | 93 | 4.89E-05 | 4.92E-05 | *Tie* |
| **SNP ID** | **MAF** | **Minor Allele Count** | **Major Allele Count** | ***P*-value** | **Mixed model**  ***P*-value** | **Gene(s)** |
| 3R_15435018_SNP | 0.3205 | 50 | 106 | 2.50E-05 | 4.94E-05 | *CG17751* |
| X_19021094_SNP | 0.1235 | 21 | 149 | 5.26E-05 | 4.95E-05 | *tgy* |
| 2L_10557230_SNP | 0.2658 | 42 | 116 | 4.10E-05 | 4.96E-05 | *Trim9* |
| 3L_1752422_SNP | 0.3252 | 53 | 110 | 4.82E-05 | 5.09E-05 | *Cht2* |
| 2R_20959318_SNP | 0.2485 | 41 | 124 | 3.01E-05 | 5.29E-05 | *gol* |
| 3R_7178120_SNP | 0.1488 | 25 | 143 | 5.23E-05 | 5.29E-05 | *pros* |
| 3L_5234682_SNP | 0.2485 | 42 | 127 | 6.06E-05 | 5.33E-05 | *shep* |
| 3L_3134029_SNP | 0.1588 | 27 | 143 | 4.88E-05 | 5.41E-05 | *CG11537* |
| X_4322495_SNP | 0.1287 | 22 | 149 | 5.02E-05 | 5.44E-05 | *bi* |
| X_10762998_SNP | 0.4235 | 72 | 98 | 4.94E-05 | 5.48E-05 | *CG2157* |
| 2R_17349447_SNP | 0.05848 | 10 | 161 | 5.90E-05 | 5.49E-05 | *Sdc* |
| 3L_6026157_SNP | 0.05882 | 10 | 160 | 4.96E-05 | 5.49E-05 | *CG10477* |
| 3L_1461925_SNP | 0.2529 | 43 | 127 | 6.23E-05 | 5.51E-05 | *Ptp61F* |
| 3L_4516376_SNP | 0.184 | 30 | 133 | 3.86E-05 | 5.53E-05 | *Tie* |
| 3L_18036228_SNP | 0.3354 | 54 | 107 | 4.65E-05 | 5.58E-05 | *Eip75B* |
| 2R_17976501_SNP | 0.3152 | 52 | 113 | 6.31E-05 | 5.62E-05 | *CG13502* |
| 2R_11756975_SNP | 0.07738 | 13 | 155 | 6.30E-05 | 5.63E-05 | *sli* |
| 2R_12113827_SNP | 0.3395 | 55 | 107 | 6.89E-05 | 5.68E-05 | *CG15706* |
| 3L_19224314_SNP | 0.1562 | 25 | 135 | 3.44E-05 | 5.72E-05 | *fz2* |
| 3L_4056180_SNP | 0.1657 | 28 | 141 | 5.68E-05 | 5.72E-05 | *CG1136* |
| 3L_5421862_SNP | 0.2 | 33 | 132 | 5.45E-05 | 5.73E-05 | *IP-delta* |
| 3L_1752429_SNP | 0.3373 | 57 | 112 | 5.74E-05 | 5.74E-05 | *Cht2* |
| 3L_6009913_SNP | 0.1479 | 25 | 144 | 5.13E-05 | 5.77E-05 | *CG32406* |
| 3L_3735843_SNP | 0.345 | 59 | 112 | 6.18E-05 | 5.83E-05 | *CG32264* |
| X_5319030_SNP | 0.4793 | 81 | 88 | 6.42E-05 | 5.83E-05 | *CG4165* |
| 2R_20831971_SNP | 0.05294 | 9 | 161 | 4.08E-05 | 5.90E-05 | *CG15861* |
| 2R_20831989_SNP | 0.05294 | 9 | 161 | 4.08E-05 | 5.90E-05 | *CG15861* |
| 2R_20832172_SNP | 0.05294 | 9 | 161 | 4.08E-05 | 5.90E-05 | *CG15861* |
| 2L_17217460_SNP | 0.125 | 20 | 140 | 6.03E-05 | 5.91E-05 | *beat-lllc* |
| 3L_2558857_SNP | 0.1325 | 22 | 144 | 6.81E-05 | 5.92E-05 | *msn* |
| 2R_13258575_SNP | 0.09884 | 17 | 155 | 5.18E-05 | 6.00E-05 | *mbl* |
| 3L_4676686_SNP | 0.05233 | 9 | 163 | 6.02E-05 | 6.00E-05 | *axo* |
| 3L_16158308_SNP | 0.1355 | 21 | 134 | 4.74E-05 | 6.07E-05 | *CG5151* |
| 2L_6347147_SNP | 0.1212 | 20 | 145 | 6.31E-05 | 6.08E-05 | *CG9497* |
| X_21217308_SNP | 0.1579 | 27 | 144 | 4.99E-05 | 6.14E-05 | *dod* |
| 3L_2622101_SNP | 0.3054 | 51 | 116 | 6.73E-05 | 6.20E-05 | *Pxn* |
| 2L_8639278_SNP | 0.08485 | 14 | 151 | 9.43E-05 | 6.23E-05 | *Sema-1a* |
| 3L_4633396_SNP | 0.1687 | 28 | 138 | 7.79E-05 | 6.27E-05 | *axo* |
| 3L_2622096_SNP | 0.3072 | 51 | 115 | 6.43E-05 | 6.29E-05 | *Pxn* |
| **SNP ID** | **MAF** | **Minor Allele Count** | **Major Allele Count** | ***P*-value** | **Mixed model**  ***P*-value** | **Gene(s)** |
| X_17662217_SNP | 0.4 | 60 | 90 | 9.68E-05 | 6.32E-05 | *unc-4* |
| 3L_4781960_SNP | 0.2353 | 40 | 130 | 6.69E-05 | 6.39E-05 | *Gef64C* |
| 3L_16149472_SNP | 0.08235 | 14 | 156 | 5.72E-05 | 6.45E-05 | *CG5151* |
| 2R_17349404_SNP | 0.05882 | 10 | 160 | 7.00E-05 | 6.51E-05 | *Sdc* |
| 2R_4513814_SNP | 0.07784 | 13 | 154 | 6.74E-05 | 6.54E-05 | *beta3GalTII* |
| 2R_6343064_SNP | 0.4785 | 78 | 85 | 5.05E-05 | 6.55E-05 | *G-oalpha47A* |
| 3L_3286773_SNP | 0.07558 | 13 | 159 | 6.46E-05 | 6.58E-05 | *CG12017* |
| X_21228530_SNP | 0.1279 | 22 | 150 | 5.00E-05 | 6.76E-05 | *sol* |
| 2L_16625000_SNP | 0.1375 | 22 | 138 | 4.18E-05 | 6.77E-05 | *CG42389* |
| 3L_5421889_SNP | 0.2321 | 39 | 129 | 4.98E-05 | 6.78E-05 | *IP-delta* |
| 3L_4342395_SNP | 0.2202 | 37 | 131 | 8.22E-05 | 6.80E-05 | *Cip4* |
| 3L_9057757_SNP | 0.2202 | 37 | 131 | 7.35E-05 | 6.82E-05 | *Argk* |
| 3L_16157916_SNP | 0.1333 | 22 | 143 | 7.70E-05 | 6.82E-05 | *CG5151* |
| 2R_21028144_SNP | 0.05294 | 9 | 161 | 6.56E-05 | 6.85E-05 | *Tkr/Lov* |
| 2R_17107726_SNP | 0.09816 | 16 | 147 | 8.43E-05 | 6.95E-05 | *ASPP* |
| 2R_20297885_SNP | 0.472 | 76 | 85 | 5.45E-05 | 7.04E-05 | *SerT* |
| 2R_6354443_SNP | 0.1975 | 32 | 130 | 7.71E-05 | 7.08E-05 | *Caf1-105* |
| 3L_1076520_SNP | 0.2917 | 49 | 119 | 5.64E-05 | 7.12E-05 | *bab1* |
| 2L_5921028_SNP | 0.3152 | 52 | 113 | 5.26E-05 | 7.14E-05 | *bchs* |
| 3L_1218317_SNP | 0.1856 | 31 | 136 | 7.56E-05 | 7.14E-05 | *mwh* |
| 3L_7450209_SNP | 0.2882 | 49 | 121 | 7.07E-05 | 7.34E-05 | *ppk26* |
| 2R_20860657_SNP | 0.05263 | 9 | 162 | 7.71E-05 | 7.45E-05 | *Phk-3* |
| 3L_16156355_SNP | 0.1325 | 22 | 144 | 6.87E-05 | 7.48E-05 | *CG5151* |
| 3L_5693934_SNP | 0.08824 | 15 | 155 | 7.19E-05 | 7.56E-05 | *sif* |
| X_21148637_SNP | 0.185 | 32 | 141 | 6.65E-05 | 7.62E-05 | *CG32521* |
| 3L_5627452_SNP | 0.05263 | 9 | 162 | 7.95E-05 | 7.62E-05 | *Blimp-1* |
| 3L_15304179_SNP | 0.2381 | 40 | 128 | 7.95E-05 | 7.66E-05 | *CG7255* |
| 3L_15304189_SNP | 0.2381 | 40 | 128 | 7.97E-05 | 7.68E-05 | *CG7255* |
| 3L_6026147_SNP | 0.05882 | 10 | 160 | 7.30E-05 | 7.83E-05 | *CG10477* |
| 3L_4640127_SNP | 0.1657 | 28 | 141 | 8.80E-05 | 7.88E-05 | *axo* |
| X_20129697_SNP | 0.05263 | 9 | 162 | 7.62E-05 | 7.90E-05 | *CG42577* |
| 2R_20818566_SNP | 0.05233 | 9 | 163 | 8.11E-05 | 7.97E-05 | *CG30427* |
| 2R_20825678_SNP | 0.05233 | 9 | 163 | 8.11E-05 | 7.97E-05 | *CG3760* |
| 2R_20836499_SNP | 0.05233 | 9 | 163 | 8.11E-05 | 7.97E-05 | *CG12851* |
| 2R_20860620_SNP | 0.05233 | 9 | 163 | 8.11E-05 | 7.97E-05 | *Phk-3* |
| 2R_20866557_SNP | 0.05233 | 9 | 163 | 8.11E-05 | 7.97E-05 | *emp* |
| 2R_20872101_SNP | 0.05233 | 9 | 163 | 8.11E-05 | 7.97E-05 | *emp* |
| 2R_20933979_SNP | 0.05233 | 9 | 163 | 8.11E-05 | 7.97E-05 | *gsb-n* |
| 2R_20966329_SNP | 0.05233 | 9 | 163 | 8.11E-05 | 7.97E-05 | *gol* |
| **SNP ID** | **MAF** | **Minor Allele Count** | **Major Allele Count** | ***P*-value** | **Mixed model**  ***P*-value** | **Gene(s)** |
| 2R_20996373_SNP | 0.05233 | 9 | 163 | 8.11E-05 | 7.97E-05 | *Tkr* |
| 2R_21018829_SNP | 0.05233 | 9 | 163 | 8.11E-05 | 7.97E-05 | *Tkr* |
| 2R_21022218_SNP | 0.05233 | 9 | 163 | 8.11E-05 | 7.97E-05 | *Tkr* |
| 3L_15943861_SNP | 0.4936 | 77 | 79 | 5.85E-05 | 8.14E-05 | *Pka-C3* |
| X_12997272_SNP | 0.5 | 83 | 83 | 6.12E-05 | 8.15E-05 | *CG43314* |
| 3L_1100091_SNP | 0.2544 | 43 | 126 | 8.96E-05 | 8.16E-05 | *bab1* |
| 2L_9336738_SNP | 0.09639 | 16 | 150 | 7.76E-05 | 8.20E-05 | *Eaat1* |
| 2R_20817924_SNP | 0.05263 | 9 | 162 | 8.72E-05 | 8.26E-05 | *CG30427* |
| 2R_20980222_SNP | 0.05263 | 9 | 162 | 8.72E-05 | 8.26E-05 | *Tkr* |
| 2R_6354251_SNP | 0.1963 | 32 | 131 | 9.00E-05 | 8.38E-05 | *Caf1-105* |
| 2R_6290625_SNP | 0.1964 | 33 | 135 | 9.24E-05 | 8.44E-05 | *SLO2* |
| 2R_6055012_SNP | 0.4717 | 75 | 84 | 7.26E-05 | 8.54E-05 | *KCNQ* |
| X_2969636_SNP | 0.08187 | 14 | 157 | 7.34E-05 | 8.56E-05 | *kirre* |
| 2R_20876342_SNP | 0.05263 | 9 | 162 | 8.75E-05 | 8.58E-05 | *CG3829* |
| 3L_8913324_SNP | 0.494 | 82 | 84 | 9.46E-05 | 8.61E-05 | *Tsp66E* |
| 3R_24243541_SNP | 0.06471 | 11 | 159 | 8.63E-05 | 8.62E-05 | *beat-VI* |
| X_10122673_SNP | 0.3526 | 61 | 112 | 8.28E-05 | 8.68E-05 | *CG34408* |
| X_21148584_SNP | 0.186 | 32 | 140 | 7.68E-05 | 8.69E-05 | *CG32521* |
| 3L_5582009_SNP | 0.09357 | 16 | 155 | 8.99E-05 | 8.70E-05 | *l(3)psg2* |
| X_13848849_SNP | 0.1607 | 27 | 141 | 8.81E-05 | 8.76E-05 | *mamo* |
| X_20423349_SNP | 0.4497 | 76 | 93 | 9.51E-05 | 8.93E-05 | *CG42267* |
| 2L_9349531_SNP | 0.09639 | 16 | 150 | 7.50E-05 | 8.98E-05 | *CG43350* |
| 2R_20935920_SNP | 0.05263 | 9 | 162 | 8.51E-05 | 9.00E-05 | *gsb-n* |
| 2R_5340696_SNP | 0.1097 | 17 | 138 | 6.95E-05 | 9.25E-05 | *Camta* |
| 3L_13250407_SNP | 0.09884 | 17 | 155 | 9.62E-05 | 9.36E-05 | *caps* |
| 3L_7269868_SNP | 0.2176 | 37 | 133 | 7.79E-05 | 9.42E-05 | *unc-13-4A* |
| 3L_13264634_SNP | 0.439 | 72 | 92 | 9.84E-05 | 9.50E-05 | *caps* |
| 2L_10853093_SNP | 0.09524 | 16 | 152 | 7.93E-05 | 9.56E-05 | *CG17086* |
| 3R_16031656_SNP | 0.05 | 8 | 152 | 6.59E-05 | 9.57E-05 | *CG34139* |
| 3L_15932321_SNP | 0.08235 | 14 | 156 | 9.48E-05 | 9.70E-05 | *Pka-C3* |
| X_13641976_SNP | 0.2831 | 47 | 119 | 9.31E-05 | 9.75E-05 | *CG10990* |
| 3L_5686074_SNP | 0.0578 | 10 | 163 | 9.53E-05 | 9.78E-05 | *sif* |
| 2L_17751052_SNP | 0.4771 | 73 | 80 | 5.40E-05 | 9.78E-05 | *CadN* |
| 2R_10634940_SNP | 0.1988 | 33 | 133 | 9.33E-05 | 9.84E-05 | *AttA* |
| 3R_7964677_SNP | 0.2308 | 36 | 120 | 4.07E-05 | 9.90E-05 | *dpr15* |
| 2R_13258724_SNP | 0.1111 | 19 | 152 | 8.77E-05 | 9.92E-05 | *mbl* |
| 2L_17764908_SNP | 0.2014 | 29 | 115 | 7.10E-05 | 9.95E-05 | *CadN2* |

**Table S3.** Type III ANOVA table for *Wolbachia* and inversion covariates

| **Factor** | **Df** | **Sum Sq** | **RSS** | **AIC** | **F value** | **Pr(>F)** |
| --- | --- | --- | --- | --- | --- | --- |
| *Wolbachia* | 1 | 4.806 | 1503.9 | 396.12 | 0.5162 | 0.4735 |
| In_2L_t | 2 | 2.409 | 1501.5 | 393.84 | 0.1293 | 0.8788 |
| In_2R_NS | 2 | 4.403 | 1503.5 | 394.07 | 0.2364 | 0.7897 |
| In_3R_P | 2 | 41.525 | 1540.6 | 398.29 | 2.2299 | 0.1109 |
| In_3R_K | 2 | 18.551 | 1517.6 | 395.69 | 0.9962 | 0.3715 |
| In_3R_Mo | 2 | 1.335 | 1500.4 | 393.72 | 0.0717 | 0.9308 |
